# The butterfly effect: amplification of local changes along the temporal processing hierarchy

**DOI:** 10.1101/102590

**Authors:** Y Yeshurun, M Nguyen, U. Hasson

## Abstract

Changing just a few words in a story can induce a substantial change in the overall narrative. How does the brain accumulate and process local and sparse changes, creating a unique situation model of the story, over the course of a real-life narrative? Recently, we mapped a hierarchy of processing timescales in the brain: from early sensory areas that integrate information over 10s-100s ms, to high-order areas that integrate information over many seconds to minutes. Based on this hierarchy, we hypothesize that early sensory areas would be sensitive to local changes in word use, but that there will be increasingly divergent neural responses along the processing hierarchy as higher-order areas accumulate and amplify these local changes. To test this hypothesis, we created two structurally related but interpretively distinct narratives by changing some individual words. We found that the neural response distance between the stories was amplified as story information is transferred from low-level regions (e.g. early auditory cortex) to high-level regions (e.g precuneus and prefrontal cortex) and that the neural difference between stories is highly correlated with an area’s ability to integrate information over time. Our results suggest a neural mechanism by which two similar situations become easy to distinguish.

## Introduction

Stories unfold over many minutes and are organized in temporarily nested structures: paragraphs are made of sentences, which are made of words, which are made of phonemes. Understanding a story therefore requires processing the story at multiple timescales such that words and phonemes are processed in a relatively short temporal window while sentences and paragraphs are processed at longer timescales. It was recently suggested that these timescales of language processing (i.e. “word level” vs “paragraph level”) are represented hierarchically along the cortical surface (Ding, Melloni, Zhang, Tian, & Poeppel, 2016; Hasson, Chen, & Honey, 2015; Kiebel, Daunizeau, & Friston, 2008; Murray et al., 2014). Previously, we defined a temporal receptive window (TRW) as the length of time in which prior information from an ongoing stimulus can affect the processing of newly arriving information. We found that early sensory areas, such as auditory cortex, have short TRWs, accumulating information over very short period of time (10s-100s milliseconds, equivalent to articulating a phoneme or word), while adjacent areas along the superior temporal sulcus have intermediate TRWs (few seconds, sufficient to integrate information at the sentence level). Areas at the top of the processing hierarchy, including the temporal parietal junction (TPJ), angular gyrus, and posterior and frontal medial cortices, have long-TRWs (many seconds to minutes), sufficient to integrate information at the paragraph and narrative levels (Hasson et al., 2015; Hasson, Yang, Vallines, Heeger, & Rubin, 2008; Honey, Thesen, et al., 2012; Lerner, Honey, Silbert, & Hasson, 2011). The neural circuits enabling long-TRW areas to accumulate and integrate information over longer periods of time are not yet been elucidated. However, recent work with both electrocorticography (ECoG) and fMRI in humans (Honey et al, 2012; Stephens et al., 2013) has suggested that the ability of an area to accumulate information may be related to its intrinsic neuronal dynamics (specifically the proportion of slow fluctuations during both rest and naturalistic stimulation).

We previously (Hasson et al., 2008) proposed that the hierarchy of TRWs is conceptually analogous to the well-established hierarchy of increasing spatial receptive fields (SRF) size in visual cortex (Dumoulin & Wandell, 2008; Grill-Spector & Malach, 2004; Smith, Singh, Williams, & Greenlee, 2001). In the visual domain, cortical areas with larger SRFs integrate and summate information from downstream areas that have smaller SRFs (Kay, Winawer, Mezer, & Wandell, 2013; Press, Brewer, Dougherty, Wade, & Wandell, 2001). As a result, small changes in the visual field, which are detected by low-level areas with small SRFs, can be amplified as they are integrated up the visual hierarchy to areas with successively larger SRFs (Hasson, Hendler, Ben Bashat, & Malach, 2001). Similarly, here we suggest that local momentary changes in the content of linguistic input in the context of a narrative (e.g. “he” vs. “she) will introduce temporally local changes in the responses of areas with short TRWs. However, such temporally local changes can affect the interpretation of a sentence (e.g. “**he** built a wall” vs. “**she** built a wall”), which unfolds over a few seconds, as well as the interpretation over the overall narrative (“they didn’t **pay** for **it**; a big surprise” vs. “they didn’t **vote** for **her**; a big surprise”), which unfolds over many minutes. Small changes may thus lead to substantial differences in the activity of high-level areas with long TRWs (Fig. 1).

**Fig. 1.**
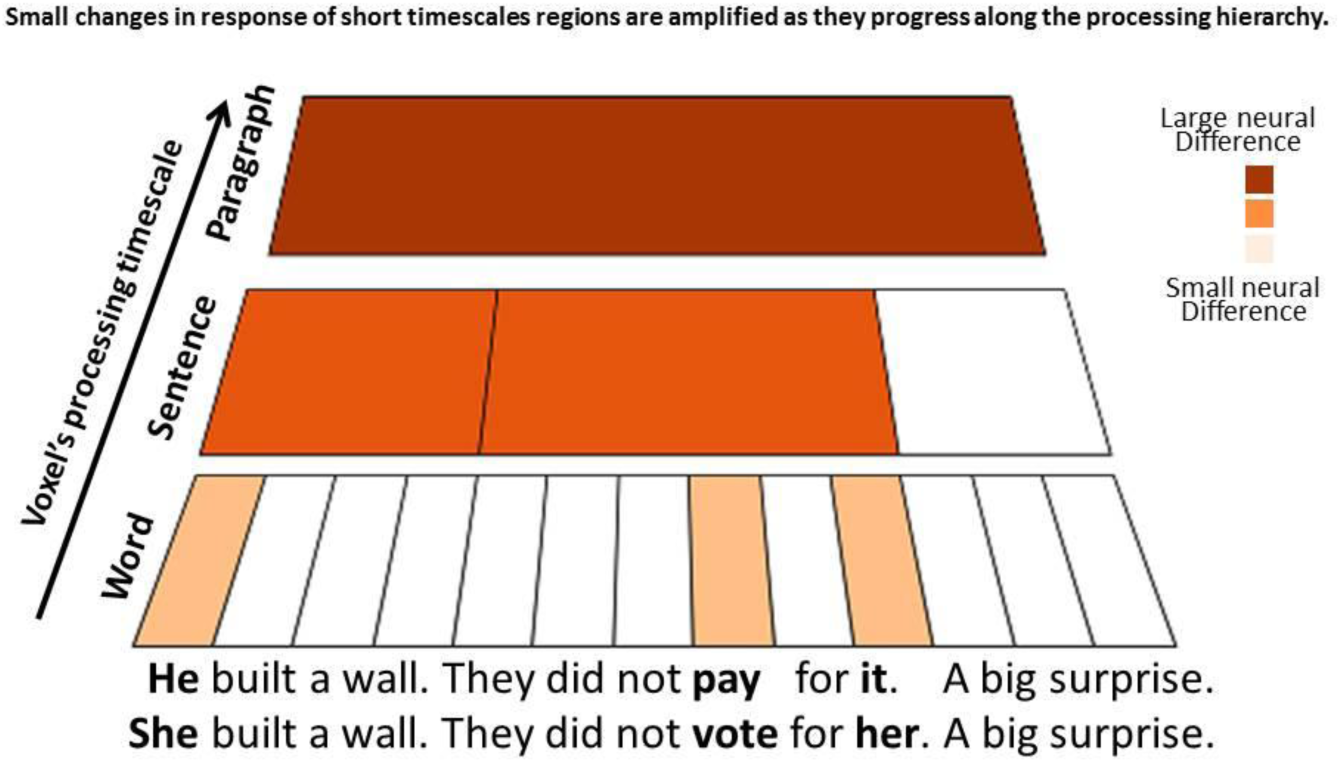
Hypothesis. Two paragraphs with three sentences each (lower panel). Only three words (out of 13) differ between paragraphs, but these small local changes result in large changes in the overall narrative. We hypothesis that voxels with short TRWs will have relatively small neural differences between narratives (bright orange), whereas voxels with long TRWs will have relatively large neural difference (brown).

In the present work, we tested the prediction that temporally local, sparse changes of words in a narrative will be accumulated and amplified along the processing timescale hierarchy. Just as small changes in word choice may lead to large changes in overall story plot, we predict that these changes will lead to increasingly divergent neural response patterns along the timescale hierarchy such that short-TRW areas will show only small differences between stories while long-TRW areas will show large changes (Fig. 1). Moreover, we predict that areas with larger neural differences between stories will have slower BOLD signal fluctuations, which enable these areas to accumulate more information over time. To test these predictions, we scanned subjects using functional magnetic resonance imaging (fMRI) while they listened to one of two stories. The two stories had the same grammatical structure, but differed in one to three words in each sentence, resulting in two distinct, yet fully coherent, narratives (Fig. 2A). To test for increasing divergence of neural responses to these two stories along the timescale hierarchy, we measured the Euclidean distance between timecourses for the two stories in each voxel. In addition, we scanned a subset of the subjects during rest, enabling us to measure intrinsic, low frequency fluctuations in the BOLD signal. In line with our hypothesis, we found a gradual divergence of the neural responses between the two stories along the timescales hierarchy. The greatest divergence occurred in areas with long TRWs and concomitant slow cortical dynamics. Our results suggest that small neural differences in low-level areas, which arise from local differences in the speech sounds, are gradually accumulated and amplified as information is transmitted from one level of the processing hierarchy to the next, ultimately resulting in distinctive neural representations for each narrative at the top of the hierarchy.

**Fig. 2.**
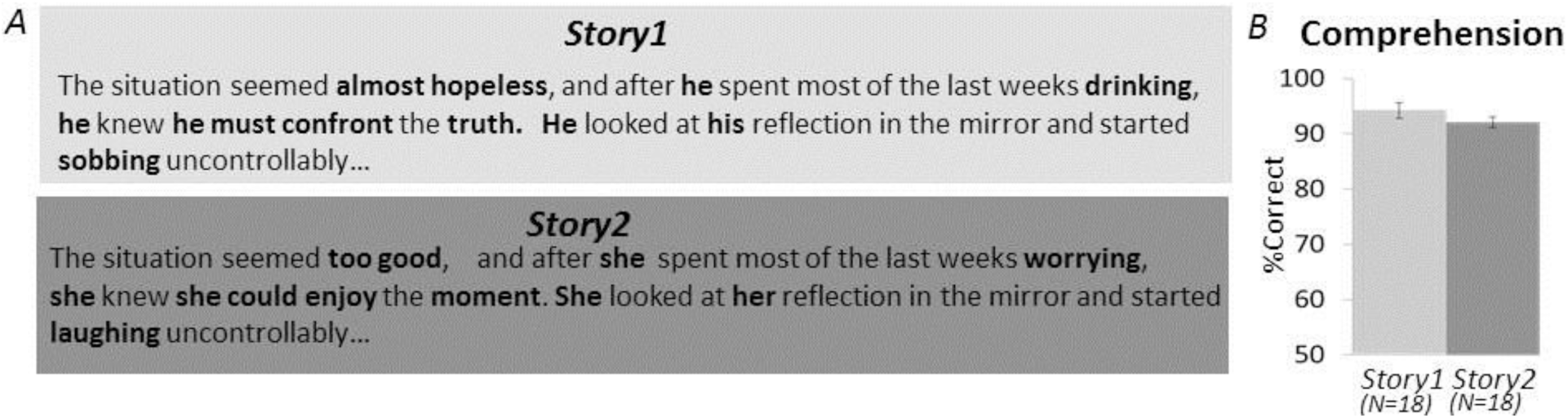
Stimuli. We scanned subjects while they listened to one of two stories that had the same grammatical structure, but differed in 33% of the words, resulting in two distinct narratives. (A) In *Story1*, a man is obsessed with his ex-girlfriend, meets a hypnotist, and then becomes fixated on Milky Way candy bars (light gray is excerpted from *Story1*). In *Story2*, a woman is obsessed with an ‘American Idol’ judge, meets a psychic, and then becomes fixated on vodka (dark gray, *Story2*). Marked in bold are words that differ between the two stories. (B) Participants’ performance on the behavioral questionnaire revealed that the comprehension level for each of the stories was high (*Story1*: 94.2±0.06%; *Story2*: 92.1±0.05%), with no difference between the two groups (t(34) = 0.59, p=0.28).

## Results

### Behavioral results: Similar comprehension of the two stories

In order to assess subjects’ comprehension of the story, we presented subjects with 28 questions immediately after the scan ended. Comprehension level for each of the stories was high (*Story1*: 94.2±0.06%; *Story2*: 92.1±0.05%), with no difference between the two stories groups (t(34) = 0.59, p=0.28), indicating that both stories were equally comprehensible (Fig.2B).

### Increased neural difference between the stories from short- to long-timescale regions

We were interested in the differences in neuronal response of subjects listening to *Story1* compared to those listening to *Story2*. In order to test for such differences, we used a Euclidean distance metric (see Methods for details). We calculated the Euclidean distance between the two groups’ response timecourses within all voxels that responded reliably to both stories (7591 voxels, see Methods). We ranked the 7591 reliable voxels based on their neural Euclidean distance, and then divided them into five equal-sized bins. These bins are presented in Fig.3A, showing bins with relatively small difference between the stories (-0.4±0.01, light orange color) up to bins with relatively large difference between the stories (3.14±0.02, brown color). Regions with relatively small difference between the stories include short-TRW areas such as auditory cortex, medial STS and ventral posterior STS (marked in light orange). Regions with relatively large difference between the stories include medium- to long-TRW areas such as the precuneus, bilateral angular gyrus, bilateral temporal poles, and medial and lateral pre-frontal cortex (marked in dark orange and brown). In each of these areas, we also measured the similarity of the response to the two stories using a between groups ISC analysis (see methods for details), to provide an intuitive sense of the magnitude of the effects. We found that the bin with the smallest Euclidean distance had the highest mean similarity between the stories (mean = 0.51 ± 0.04). The range of ISC values among the voxels in this bin was skewed towards greater between-story similarity (range: 0.39-0.73, Fig. 3B). In contrast, the bin with the largest Euclidean distance showed very little cross-story ISC similarity (mean = 0.35 ± 0.06), with skewed probability toward areas with close to zero similarity in response patterns across the two stories (range: 0.066-0.52; Fig.3B). These differences between the bins were highly significant as revealed by 1-way ANOVA on the normalized ISC (F(1,7585) = 2622.58, p<10^-10^; Scheffe’s post-hoc comparisons revealed that all the bins’ normalized ISC significantly differ from each other).

**Fig. 3.**
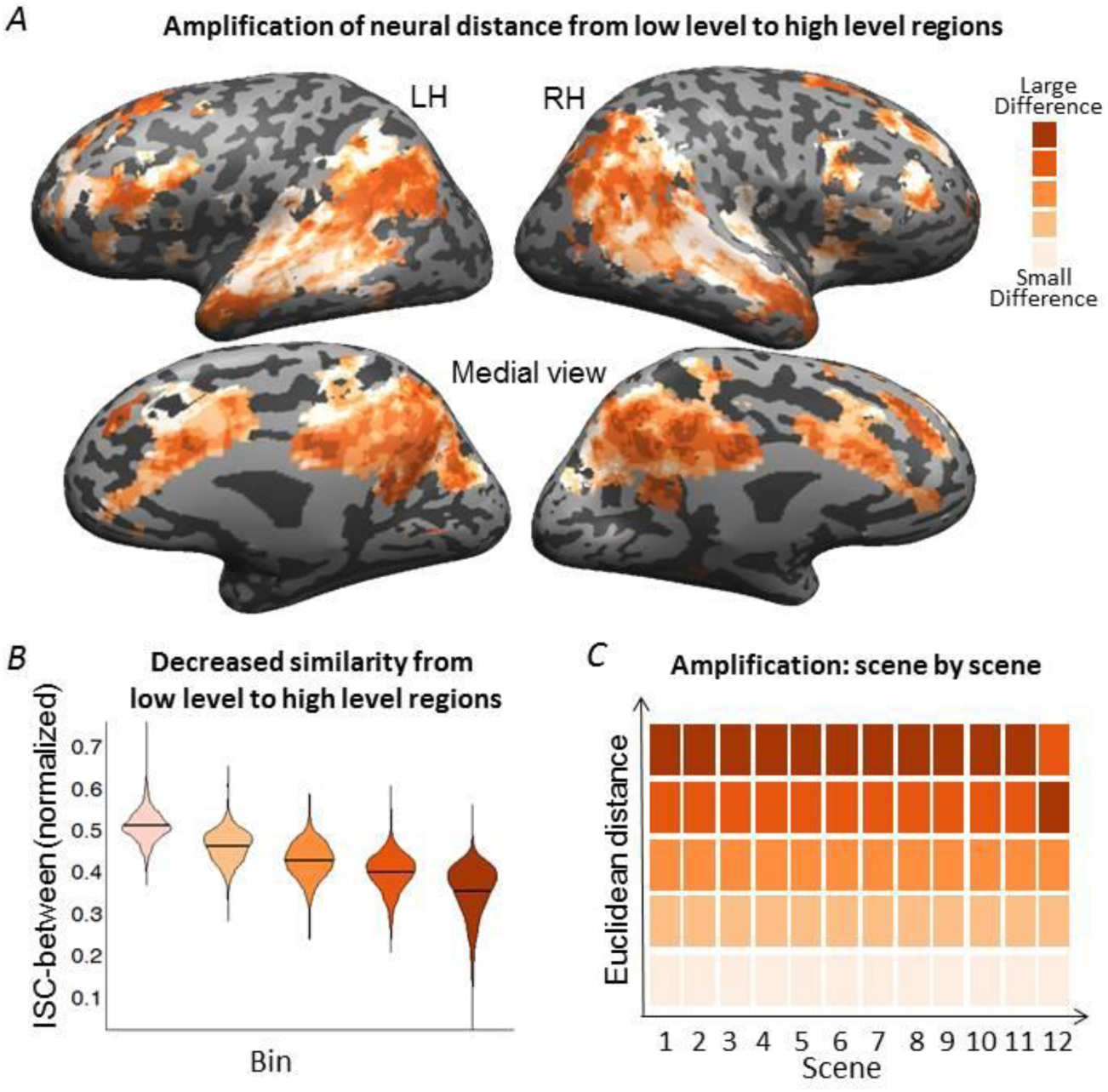
Neural results demonstrating amplification of the neural distance. (A) We calculated the Euclidean distance between the two groups’ response timecourses and ranked the voxels based on their neural Euclidean distance, and then divided them into five equal-sized bins. Small neural differences are primarily observed in and around primary auditory cortex while increasingly large neural differences are observed extending towards TPJ, precuneus, and frontal areas. (B) Normalized ISC in each of the five bins. The bin with the smallest Euclidean distance (marked in light orange) showed the highest ISC, whereas the bin with the largest Euclidean distance showed the lowest ISC between the stories (brown color). These differences between the bins were highly significant. (C) We calculated the Euclidean distance between the stories’ neural response in each of the 12 scenes, and then rank ordered these average distance values. In 11 of the 12 scenes, the ordering of bins was the same as for the overall story, with only small change in the Euclidean distance between the areas with large differences in the last segment.

## Amplification pattern is replicated in 11 out of 12 scenes of the story

We next tested whether the amplification of neural differences from short- to long-TRW areas occurred in every scene of the story. To that end, we calculated the Euclidean distance between the stories’ neural response in each of the 12 scenes, averaged them across voxel bins (defined over the entire story), and then rank ordered these average distance values. In 11 of the 12 scenes, the ordering of bins was the same as for the overall story, with only small change in the Euclidean distance between the areas with large differences in the last segment (Fig.3C). The consistency of ordering across the 12 independent segments suggests that the amplification of neural distance from low-level regions to high-level regions is robust and stable.

### Significant correlation between neural distance and capacity to accumulate information over time

Finally, we asked whether the difference in neural response to the two stories was related to the capacity to accumulate information over time. To characterize this capacity, we calculate a time receptive window (TRW) index using an independent data set obtained while subjects listened to an intact and word scrambled versions of another story (see methods for details). Naked-eye comparison of the TRW index brain map (Fig.4A, upper panel) and the two stories Euclidean distance map (Fig.4A, lower panel) revealed high similarity between the maps. Indeed, we found, at a voxel-by-voxel basis, that the larger the difference in the neural responses between the stories, the larger the voxel’s TRW index (r=0.426, p<0.001) (Fig.4B). As an area’s TRW size may be related to intrinsically slower cortical dynamics in these areas (Honey et al., 2012; Stephens et al., 2013), we also calculated the proportion of low frequency power during a resting state scan. We found that voxels with larger neural difference between stories also had greater proportion of low frequency power (r=0.246, p<0.001) (Fig. 4C).

**Fig. 4.**
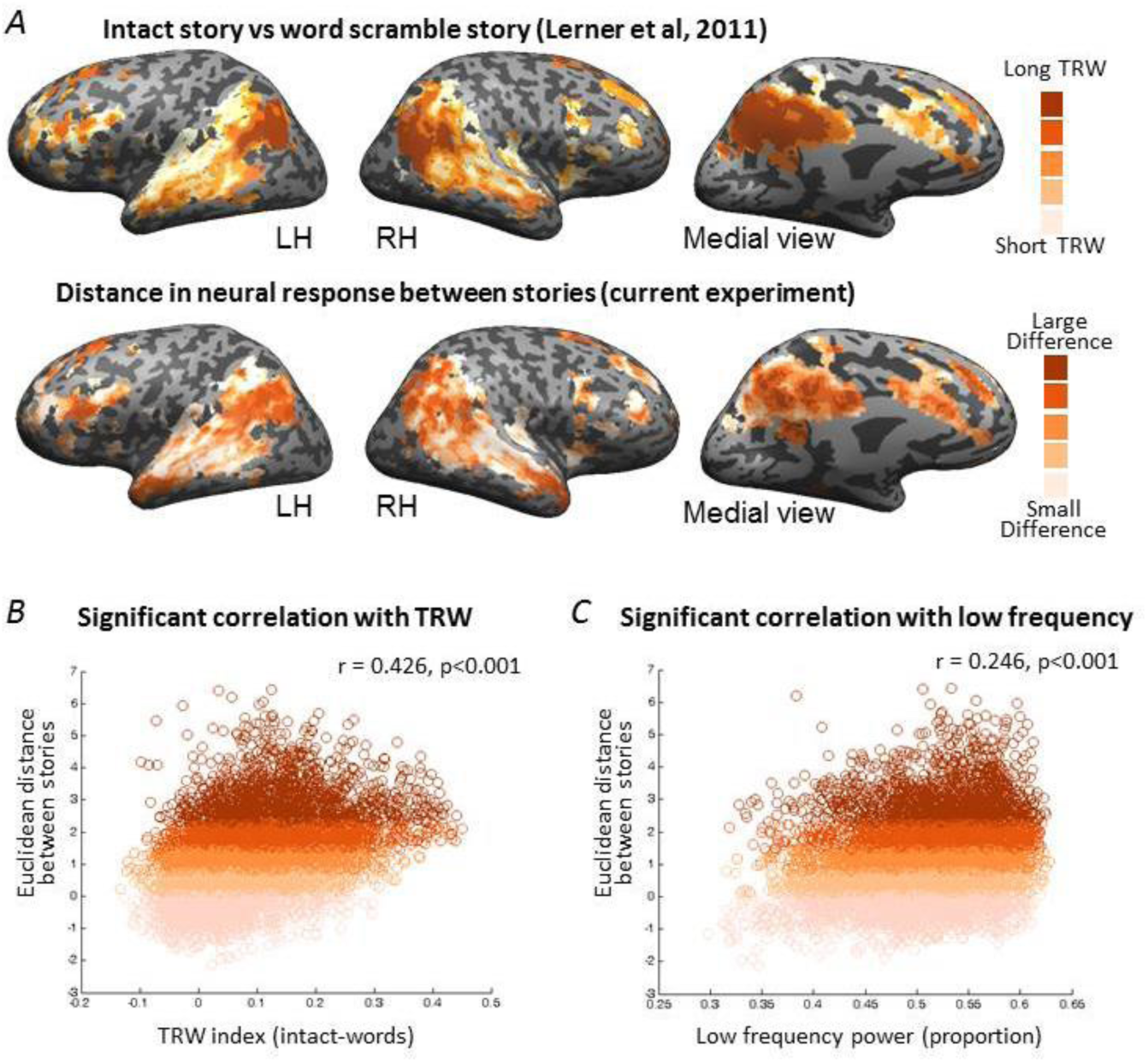
Correlation with processing timescales. (A) TRW index brain map (upper panel) and the two stories Euclidean distance map (lower panel). (B) Scatter plot of the voxel’s TRW-index and the Euclidean distance between the stories. The larger the voxel’s TRW index, the larger the difference in the neural responses between the stories (C) Scatter plot of the voxel’s proportion of low frequency power during a resting state scan and the Euclidean distance between the stories. Voxels with greater low frequency power proportion was correlated with larger neural difference between stories.

## Discussion

Small, local changes of word choice in a story can completely alter the narrative. In this study, we predicted that the accumulation and amplification of sparse, temporally local word changes will be reflected in increasing divergent neural response along the timescale processing hierarchy. In line with our predictions, we found that short timescale areas, including primary auditory cortex, showed only small neural differences in response to local word changes. This finding is consistent with observations that these early auditory areas process transient and rapidly changing sensory input (Okada et al., 2010; Poeppel, 2003), such that brief local alternations in the sound structure will only induce brief and local alternations in the neural responses across the two stories. However, just as these small word changes are amplified in the overall interpretation of the whole story, we found that small neural differences in early sensory areas were accumulated and amplified as they traversed the processing timescale hierarchy: while short-TRW areas showed small neural differences (0.62 normalized correlation between stories), long-TRWs areas, including TPJ, angular gyrus, PCC, and dmPFC, showed large neural differences (0.16 normalized correlation between stories) (Fig. 3). Moreover, the difference in neural response between stories was significantly correlated with both TRW length (r = .426, p<.001) and slower cortical dynamics (r = .14, p<.02).

The topographical hierarchy of processing timescales along the cortical surface (Hasson et al., 2015) is supported by single unit analysis (Murray et al., 2014), ECoG analysis (Honey, Thesen, et al., 2012), fMRI analysis (Baldassano et al., 2016; Hasson et al., 2008; Lerner et al., 2011), MEG analysis (Ding et al., 2016), computational models (Kiebel et al., 2008), and resting state functional connectivity (Margulies et al., 2016; Sepulcre, Sabuncu, Yeo, Liu, & Johnson, 2012). Based on this hierarchy, it was hypothesized that high-level cortical areas encode slowly changing states of the world, while low-level areas encode fast changes (Honey, Thesen, et al., 2012; Kiebel et al., 2008; Stephens, Honey, & Hasson, 2013). In line with this hypothesis, we found that low-level areas showed relatively similar neural responses to the two stories, consistent with the strong similarity of the stories at the word and acoustic level. However, the neural responses in high-level areas, which integrate information over longer periods of time, were very different for the two stories, consistent with the strong dissimilarity of the stories at the narrative and situation model level. These results are in line with a very recent study demonstrating that high-level regions have a more stable neural state, which slowly changes at large event boundaries in the narrative (Baldassano et al., 2016). We suggest that areas with long processing timescales gradually construct and retrain a situation model (van Dijk & Kintsch, 1983; Zwaan & Radvansky, 1998) of the event structure based on the information which is gathered as the story unfolds over time (Speer, Zacks, & Reynolds, 2007; Zacks et al., 2001).

The topography of timescale processing is consistent with the previously proposed linguistic hierarchies: low-level regions (A1+) represent phonemes (Arsenault & Buchsbaum, 2015; Humphries, Sabri, Lewis, & Liebenthal, 2014), syllables (Evans & Davis, 2015) and pseudowords (Binder et al., 2000), while medium-level regions (areas along A1+ to STS) represent sentences (Fedorenko et al., 2016; Pallier, Devauchelle, & Dehaene, 2011). At the top of the hierarchy, high-level regions (bilateral TPJ, precuneus, mPFC) represent paragraphs up to the whole narrative (Hickok & Poeppel, 2007; Lerner et al., 2011; Price, Bonner, Peelle, & Grossman, 2015). In these studies, the different parts of the processing hierarchy were characterized by comparing neural responses to meaningful versus less meaningful stimuli (e.g. words vs pseudowords, sentences vs scrambled sentences). Here, we show how such a topographical hierarchy allows for the accumulation and integration of local changes needed for the processing of two coherent, yet markedly different narratives, which are told using the exact same grammatical structure.

Understanding a narrative requires more than just understanding the individual words in the narrative. For example, children with hydrocephaly, a neurodevelopmental disorder that is associated with brain anomalies in regions including the posterior cortex, have well-developed word decoding but concomitant poor understanding of narrative constructed from the same words (Barnes & Dennis, 1992; Barnes, Faulkner, & Dennis, 2001). In Parkinson’s Disease (PD) and early Alzheimer’s Disease (AD), researchers have found that individual word comprehension is relatively intact, yet the ability to infer the meaning of the text is impaired (Chapman, Anand, Sparks, & M., 2006). In PD, this deficit in the organization and interpretation of narrative discourse was associated with reduced cortical volume in several brain regions, including the superior part of the left STS and the anterior cingulate (Ash et al., 2011). Our finding that these regions demonstrated large differences between the stories (Fig. 3) suggests that reduced volume of cortical circuits with long processing timescales may result in reduced capability to understand temporally extended narratives.

The phenomenon we described here, in which local changes in the input generate a large change in the meaning, is ubiquitous: a small change in eye gaze differentiates happiness from anger; a small change in intonation differentiates comradery from mockery; a small change in hand pressure differentiates comfort from threat. What is the neural mechanism underlying this phenomenon? In vision, researchers have shown that low-level visual areas with small spatial receptive fields (SRFs) are sensitive to spatially confined changes in the visual field, but that these small changes are accumulated and integrated along the visual processing stream such that high-level areas, with large SRFs can show substantively different neural responses from a very small change in visual stimuli. For example, it was shown that the fusiform face area is more sensitive to holistic changes in the picture (vase vs face) than to local features and it was suggested that the fusiform face area, which is a high-order visual region, spatially group many local features to create the holistic percept (Hasson et al., 2001). Analogously, our results suggest that regions which are capable to group information across many different time points (i.e. regions with long processing timescales) are highly sensitive to holistic changes (i.e. at the narrative level). To our knowledge, this is the first study to systematically map the gradual amplification of temporally confined changes along the processing timescales hierarchy. Our study suggests a neural mechanism by which two similar situations become easy to distinguish.

## Methods

### Subjects

Thirty six right-handed subjects (ages 21.1±3.7) participated in the study. Eighteen subjects (9 female) heard *Story1*, and eighteen other subjects (9 female) heard *Story2*. Two subjects were discarded from the analysis due to head motion (>2 mm). This sample size of 18 subjects for each story was chosen based on previous studies in our lab that tested for similarities and differences in neural responses to naturalistic stimuli (Ames, Honey, Chow, Todorov, & Hasson, 2015; Honey, Thompson, Lerner, & Hasson, 2012; Lerner et al., 2011) as well as power analyses (Pajula & Tohka, 2016). Experimental procedures were approved by the Princeton University Committee on Activities Involving Human Subjects. All subjects provided written informed consent.

### Stimuli and Experimental Design

In the MRI scanner, participants listened to one of two stories. While the two stories had the exact same grammatical structure, they differed in one to three words per sentence, creating two distinct narratives (mean words change in a sentence = 2.47±1.7, 34.13%±20 of the words in the sentence). In *Story1*, a man is obsessed with his ex-girlfriend, meets a hypnotist, and then becomes fixated on Milky Way candy bars (Fig. 2A, negative to positive story arc). In *Story2*, a woman is obsessed with an ‘American Idol’ judge, meets a psychic, and then becomes fixated on vodka (Fig. 2A, positive to negative story arc). The two stories were read and recorded by the same actor. The beginning of each sentence was aligned post-recording. Each story was 6:44 minutes, and was preceded by 18 s of neutral music and 3 s of silence. The story was followed by an additional 15 s of silence. These music and silence periods were discarded from all analyses.

In addition, 26 of the subjects (13 from each group) underwent a 10-min resting state scan. Subjects were instructed to stay awake, look at a gray screen, and “think on whatever they like” during the scan.

### Behavioral assessment

Immediately following scanning, each participant’s comprehension of the story was assessed using a questionnaire presented on a computer. Twenty-eight 2-forced choice questions were presented. Two-tailed Student t-tests (α = 0.05) on the forced-choice answers were conducted between the two story groups to evaluate the difference in participants’ comprehension.

### MRI Acquisition

Subjects were scanned in a 3T full-body MRI scanner (Skyra, Siemens) with a 12-channel head coil. For functional scans, images were acquired using a T2*-weighted echo planar imaging (EPI) pulse sequence [repetition time (TR), 1500 ms; echo time (TE), 28 ms; flip angle, 64°], each volume comprising 27 slices of 4 mm thickness with 0 mm gap; slice acquisition order was interleaved. In-plane resolution was 3 × 3 mm^2^ [field of view (FOV), 192 × 192 mm^2^]. Anatomical images were acquired using a T1-weighted magnetization-prepared rapid-acquisition gradient echo (MPRAGE) pulse sequence (TR, 2300 ms; TE, 3.08 ms; flip angle 9°; 0.89 mm^3^ resolution; FOV, 256 mm^2^). To minimize head movement, subjects' heads were stabilized with foam padding. Stimuli were presented using the Psychtoolbox version 3.0.10 (Pelli, 1997). Subjects were provided with MRI compatible in-ear mono earbuds (Sensimetrics model S14), which provided the same audio input to each ear. MRI-safe passive noise-canceling headphones were placed over the earbuds for noise reduction and safety.

### Data analysis Imaging analysis Preprocessing

fMRI data were reconstructed and analyzed with the BrainVoyager QX software package (Brain Innovation) and in-house software written in MATLAB (MathWorks). Preprocessing of functional scans included intra-session 3D motion correction, slice-time correction, linear trend removal, and high-pass filtering (two cycles per condition). Spatial smoothing was applied using a Gaussian filter of 6 mm full-width at half-maximum value. The complete functional dataset was transformed to 3D Talairach space (Talairach & Tournoux, 1988).

### Euclidean distance measure

We were interested in the differences in neuronal response of subjects listening to *Story1* compared to those listening to *Story2*. In order to test for such differences, we used a Euclidean distance metric in all the voxels that reliably respond to both stories (7591 voxels with Inter-subject-correlation > 0.12, see (Lerner et al., 2011) for details). In each voxel, we calculated the mean response of the 18 subjects presented with *Story1* and the mean response of the 18 subjects presented with *Story2*. This averaging resulted in two mean timecourses, one for *Story1* (S1) and one for *Story2* (S2), each with 269 time points. Next, we calculated the Euclidean distance between the timecourses S1 and S2:

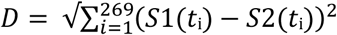

where *S*1(*t*_i_) is the mean BOLD timecourse measured in *Story1*, and *S*2(*t*_i_) is the mean BOLD timecourse measured in *Story2*. This procedure was repeated for each voxel, resulting in a distance value for each of the 7591 voxels.

In order to account for differences that arise from irrelevant sources (i.e. signal to noise ratio), we normalized Euclidian distance measures with the mean and standard deviation from null distributions generated through label shuffling. In this procedure, the two story groups were randomly shuffled such that two new pseudo-group were created, each with 9 *Story1* and 9 *Story2* timecourses. We calculated the Euclidean distance between the resulting mean responses, 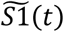 and 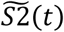, in the pseudo groups: 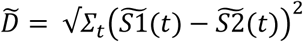. The procedure of label shuffling and computing a surrogate Euclidean distance value was repeated 50,000 times, generating a null distribution of 50,000 distance values for each voxel. We then normalized the difference in each voxel according to its specific mean and standard deviation of the null distribution: D_norm_ = (D – μ(null))/ơ(null). Reliable voxels were then ranked based on their normalized Euclidean distance value (D_norm_) and divided into five equal-sized bins. These bin categories (small to large neural difference) were then projected onto the cortical surface for visualization.

Finally, we tested whether the order of these neural difference bins was consistent across sections of the story. Do voxels with small or large neural differences over the entire story also have respectively small and large neural difference for every section of the story? Thus, we calculated D_norm_ for each of twelve scenes of the story across all the included voxels. These voxels were then sorted based on their bin from calculating D_norm_ across the entire story. The D_norm_ values for each bin were then averaged across voxels for each scene, and then rank ordered.

### ISC between the stories

The normalized Euclidean distance measure of similarity has an arbitrary scale and thus does not provide an intuitive sense of the magnitude of effects. As a complementary measure, we additionally calculated similarity using intersubject correlation (ISC, Hasson et al., 2004), which measures the degree to which neural responses to naturalistic stimuli are shared across subjects in the two different story groups. To calculate the ISC between the two story groups, for each reliable voxel, we correlated each story subject’s timeseries with the average timeseries across all of the subjects that listened to the other story. We than averaged these 36 correlation values to get an estimation of the similarity of the responses between the stories in each voxel (ISC_b_). Finally, we calculated ISC within each story (ISC_w1_ for Story1 and ISC_w2_ for Story2) by taking the average correlation between each subject and the average of all other subjects in the same group. We used this measure of within-group similarity to normalize the between-group ISC: ISC_norm_= ISC_b_/( ISC_w1_+ ISC_w2_).

### Correlation between neural distance and capacity to accumulate information over time

Next, we calculated an index of each voxel’s temporal receptive window (TRW), i.e. the capacity to accumulate information over time, in order to test the relationship between a voxel’s processing timescale and its neural distance measure. In a previously collected data set, subjects were scanned listening to both an intact version and word-scrambled version of a story (“Pieman” by Jim O’Grady, see Lerner et al., 2011 for details). Response similarity to the intact and scrambled stories was measured using ISC as described above, resulting in a value ISC_intact_ and ISC_scram_ for each voxel. Using this independent data set, we defined the TRW index of each voxel as the difference in neural activity between the intact story and the word scrambled story: TRW = ISC_intact_ – ISC_scram_. We then calculated the correlation between neural response (D_norm_, described above) with the TRW index: r = correlation(D_norm_, TRW index). Statistical significance of this correlation coefficient (that the correlation is not zero) was computed using a Student's t distribution for a transformation of the correlation.

### Relation between neural distance and timescale of BOLD signal dynamics

Previous work has suggested that long-TRW areas have intrinsically slower neural dynamics compared to short-TRW areas, and that these slower cortical dynamics may be related to an area’s ability to accumulate information over time (Honey, Thesen, et al., 2012; Stephens et al., 2013). Based on such observations, we evaluated the neural dynamics of our reliable voxels during a resting state scan in 26 of our subjects. Following Stephens et al., 2013, we estimated the power spectra of each voxel using fast Fourier transform. We then quantified the proportion of low frequency power as the accumulating power below a fixed threshold of 0.04 Hz: 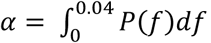, where *P*(*f*) is the power estimated as above. Finally, we calculated the Pearson correlation between proportion of low frequency power (α) and the neural response difference (D_norm_, described above) voxel by voxel. Statistical significance of this correlation coefficient (that the correlation is not zero) was computed using a Student's t distribution for a transformation of the correlation.

## Acknowledgements

### Author contributions

Y. Yeshurun and U. Hasson developed the study concept, and contributed to the study design. Data collection was performed by Y.Yeshurun. Data analysis was performed by Y. Yeshurun and M. Nguyen. Y. Yeshurun and U. Hasson drafted the manuscript, and M. Nguyen provided critical revisions. All authors approved the final version of the manuscript for submission.

## Acknowledgments

We would like to thank Gideon Dishon for his help in creating experimental materials, Prof. Anat Ninio for her help in the analysis of the stimuli grammatical structure, and Amy Price for helpful comments on the paper. This work was supported by NIH grant RO1-MH094480 (U.H, M.N and Y.Y), the Rothschild Foundation (Y.Y), and The Israel National Postdoctoral Program for Advancing Women in Science (Y.Y).

## References

Ames, D. L., Honey, C. J., Chow, M. A., Todorov, A., & Hasson, U. (2015). Contextual alignment of cognitive and neural dynamics. J Cogn Neurosci, 27(4), 655-664.

Arsenault, J. S., & Buchsbaum, B. R. (2015). Distributed neural representations of phonological features during speech perception. J Neurosci, 35(2), 634-642.

Ash, S., McMillan, C., Gross, R. G., Cook, P., Morgan, B., Boller, A., et al (2011). The organization of narrative discourse in Lewy body spectrum disorder. Brain Lang, 119(1), 30-41.

Baldassano, C., Chen, J., Zadbood, A., Pillow, J., Hasson, U., & Norman, K. (2016). Discovering event structure in continuous narrative perception and memory. bioRxiv.

Barnes, M. A., & Dennis, M. (1992). Reading in children and adolescents after early onset hydrocephalus and in normally developing age peers: phonological analysis, word recognition, word comprehension, and passage comprehension skill. J Pediatr Psychol, 17(4), 445-465.

Barnes, M. A., Faulkner, H. J., & Dennis, M. (2001). Poor reading comprehension despite fast word decoding in children with hydrocephalus. Brain Lang, 76(1), 35-44.

Binder, J. R., Frost, J. A., Hammeke, T. A., Bellgowan, P. S., Springer, J. A., Kaufman, J. N., et al (2000). Human temporal lobe activation by speech and nonspeech sounds. Cereb Cortex, 10(5), 512-528.

Chapman, S. B., Anand, R., Sparks, G., & M., C. C. (2006). Gist Distinctions in Healthy Cognitive Aging Versus Mild Alzheimer's Disease. Brain Impairment, 7(03), 223-233.

Ding, N., Melloni, L., Zhang, H., Tian, X., & Poeppel, D. (2016). Cortical tracking of hierarchical linguistic structures in connected speech. Nat Neurosci, 19(1), 158-164.

Dumoulin, S. O., & Wandell, B. A. (2008). Population receptive field estimates in human visual cortex. Neuroimage, 39(2), 647-660.

Evans, S., & Davis, M. H. (2015). Hierarchical Organization of Auditory and Motor Representations in Speech Perception: Evidence from Searchlight Similarity Analysis. Cereb Cortex, 25(12), 4772-4788.

Fedorenko, E., Scott, T. L., Brunner, P., Coon, W. G., Pritchett, B., Schalk, G., et al (2016). Neural correlate of the construction of sentence meaning. Proc Natl Acad Sci U S A, 113(41), E6256-E6262.

Grill-Spector, K., & Malach, R. (2004). The human visual cortex. Annu Rev Neurosci, 27, 649-677.

Hasson, U., Chen, J., & Honey, C. J. (2015). Hierarchical process memory: memory as an integral component of information processing. Trends Cogn Sci, 19(6), 304-313.

Hasson, U., Hendler, T., Ben Bashat, D., & Malach, R. (2001). Vase or face? A neural correlate of shape-selective grouping processes in the human brain. J Cogn Neurosci, 13(6), 744-753.

Hasson, U., Yang, E., Vallines, I., Heeger, D. J., & Rubin, N. (2008). A hierarchy of temporal receptive windows in human cortex. J Neurosci, 28(10), 2539-2550.

Hickok, G., & Poeppel, D. (2007). The cortical organization of speech processing. Nat Rev Neurosci, 8(5), 393-402.

Honey, C. J., Thesen, T., Donner, T. H., Silbert, L. J., Carlson, C. E., Devinsky, O., et al (2012). Slow cortical dynamics and the accumulation of information over long timescales. Neuron, 76(2), 423-434.

Honey, C. J., Thompson, C. R., Lerner, Y., & Hasson, U. (2012). Not lost in translation: neural responses shared across languages. J Neurosci, 32(44), 15277-15283.

Humphries, C., Sabri, M., Lewis, K., & Liebenthal, E. (2014). Hierarchical organization of speech perception in human auditory cortex. Front Neurosci, 8, 406.

Kay, K. N., Winawer, J., Mezer, A., & Wandell, B. A. (2013). Compressive spatial summation in human visual cortex. J Neurophysiol, 110(2), 481-494.

Kiebel, S. J., Daunizeau, J., & Friston, K. J. (2008). A hierarchy of time-scales and the brain. PLoS Comput Biol, 4(11), e1000209.

Lerner, Y., Honey, C. J., Silbert, L. J., & Hasson, U. (2011). Topographic mapping of a hierarchy of temporal receptive windows using a narrated story. J Neurosci, 31(8), 2906-2915.

Margulies, D. S., Ghosh, S. S., Goulas, A., Falkiewicz, M., Huntenburg, J. M., Langs, G., et al (2016). Situating the default-mode network along a principal gradient of macroscale cortical organization. Proc Natl Acad Sci U S A, 113(44), 12574-12579.

Murray, J. D., Bernacchia, A., Freedman, D. J., Romo, R., Wallis, J. D., Cai, X., et al (2014). A hierarchy of intrinsic timescales across primate cortex. Nat Neurosci, 17(12), 1661-1663.

Okada, K., Rong, F., Venezia, J., Matchin, W., Hsieh, I. H., Saberi, K., et al (2010). Hierarchical organization of human auditory cortex: evidence from acoustic invariance in the response to intelligible speech. Cereb Cortex, 20(10), 2486-2495.

Pajula, J., & Tohka, J. (2016). How Many Is Enough? Effect of Sample Size in Inter-Subject Correlation Analysis of fMRI. Comput Intell Neurosci, 2016, 2094601.

Pallier, C., Devauchelle, A. D., & Dehaene, S. (2011). Cortical representation of the constituent structure of sentences. Proc Natl Acad Sci U S A, 108(6), 2522-2527.

Pelli, D. G. (1997). The VideoToolbox software for visual psychophysics: transforming numbers into movies. Spat Vis, 10(4), 437-442.

Poeppel, D. (2003). The analysis of speech in different temporal integration windows: cerebral lateralization as 'asymmetric sampling in time'. Speech Communication, 41, 245-255.

Press, W. A., Brewer, A. A., Dougherty, R. F., Wade, A. R., & Wandell, B. A. (2001). Visual areas and spatial summation in human visual cortex. Vision Res, 41(10-11), 1321-1332.

Price, A. R., Bonner, M. F., Peelle, J. E., & Grossman, M. (2015). Converging evidence for the neuroanatomic basis of combinatorial semantics in the angular gyrus. J Neurosci, 35(7), 3276-3284.

Sepulcre, J., Sabuncu, M. R., Yeo, T. B., Liu, H., & Johnson, K. A. (2012). Stepwise connectivity of the modal cortex reveals the multimodal organization of the human brain. J Neurosci, 32(31), 10649-10661.

Smith, A. T., Singh, K. D., Williams, A. L., & Greenlee, M. W. (2001). Estimating receptive field size from fMRI data in human striate and extrastriate visual cortex. Cereb Cortex, 11(12), 1182-1190.

Speer, N. K., Zacks, J. M., & Reynolds, J. R. (2007). Human brain activity time-locked to narrative event boundaries. Psychol Sci, 18(5), 449-455.

Stephens, G. J., Honey, C. J., & Hasson, U. (2013). A place for time: the spatiotemporal structure of neural dynamics during natural audition. J Neurophysiol, 110(9), 2019-2026.

Talairach, J., & Tournoux, P. (1988). Co-planar stereotaxic atlas of the human brain: Thieme Medical Publishers, New York.

van Dijk, T. A., & Kintsch, W. (1983). Strategies of Discourse Comprehension.: Academic Press, New York.

Zacks, J. M., Braver, T. S., Sheridan, M. A., Donaldson, D. I., Snyder, A. Z., Ollinger, J. M., et al (2001). Human brain activity time-locked to perceptual event boundaries. Nat Neurosci, 4(6), 651-655.

Zwaan, R. A., & Radvansky, G. A. (1998). Situation models in language comprehension and memory. Psychological Bulletin, 123(2), 162-185.

